# Change of information by positive frequency-dependent selection in two very different models (laser-like and chirality of shell-coiling in the snail *Partula suturalis*)

**DOI:** 10.1101/2021.02.15.431340

**Authors:** W. A. Tiefenbrunner

## Abstract

Although according to the second law of thermodynamics the world tends toward maximum disorder, over millions of years evolution has given rise to an enormous variety of complex organisms. To explain this, one must assume that natural selection is a process of information acquisition. Since some years an information theory of selection exists that can quantify this change and thus helps to understand the apparent contradiction between the existence of biological complexity and the tendency toward disorder that generally prevails in nature. Here I apply this theory to examples of frequency-dependent selection (this means: in which phenotype frequency determines its fitness).

The snail *Partula suturalis* gave an evolutionary and ecologically unique and hence very valuable example of this type of selection before it became extinct about thirty years ago on its native island. Spatially separated populations with left- and right-coiled shells occurred on Moorea, but also hybridization zones. Since both types of shells were the same except for chirality, the question is whether selection happened at all. The inheritance of this character is monogenic and in this respect simple, but is complicated by the fact that it is the maternal genotype, not the own, that determines the phenotype. This causes that for the calculation of the information change by selection not the genotype or phenotype frequencies are sufficient, but one must consider their combination. The simulation shows that frequency-dependent selection in *P. suturalis* indeed increased information.

It has already been shown that selection can also be important outside animate nature, for example in the generation of laser light, which has extraordinary properties: it is monochromatic, monoaxial and monophasic. Phase selection is frequency(=density)-dependent and therefore of interest here. In selection theory the mean fitness ω is of special significance. In a laser-like model, in modeling phase selection, we find that ω=1+A^2^, where A^2^ is the the light intensity or the square of the amplitude, respectively. During selection, ω increases and, in parallel, since selection is a process of information acquisition, so does the information. Because of the connection between ω and A^2^ this also means for the laser-like model that – assuming a constant number of photons – a larger amplitude always means more information (less entropy).

## Introduction

The theory of evolution^3^, which today represents the unifying principle of biology, replaced the rigid concept of species constancy with a much more dynamic picture of animate nature. In 1858, however, Darwin and Wallace^2^ also provided an explanation of how, in the course of millions of years, organisms of incredible complexity and diversity could arise. According to this explanation, variants that are better adapted to the environment are favored by a selection process and eventually displace others. Around the same time, Maxwell^12^ and Boltzmann^1^ came to the conclusion that entropy increase means that ‘the world’ evolves towards maximum disorder. The Second Law of Thermodynamics is thus a law of increase of disorder. Due to the claim of physics that its laws should be valid for the whole, i. e. also the animate, nature, a contradiction had thus arisen, which many scientists found disturbing.

With the formulation of non-equilibrium thermodynamics, the emergence of complexity in a world where the tendency to disorder prevails was still puzzling, but at least no longer an impossibility^13^. Further progress was made by theoretical considerations concerning the origin of life^4^, for here thermodynamics and the theory of evolution inevitably meet. Evolution, it turned out, can also be of significance at the molecular level, provided auto-catalytic reactions are involved. In the context of a new science, Synergetics^9,10^, it was shown that selection even plays a role in some areas of physics. Thus, from these sciences a bridge-building had been started. Of course, a bridge is best built from both sides. The foundations for this endeavor were population genetics^8^ on the one hand and information theory^14^ on the other. Henceforth it was possible to understand selection as a process of information acquisition^15^.

R. A. Fisher 1930^8^ united genetics and evolution theory and clarified thereby also in a radical way an accusation with which already Darwin was confronted and which has to do with the variant favored by the environment. Usually the fitness of an individual is not measurable, which is why soon after the publication of Darwin’s seminal book ‘On the Origin of Species’^3^ the suspicion arose that behind the expression “survival of the fittest” there was nothing more than circular reasoning: Who is it that survives? The fittest. Who is the fittest? The one who survives. Fisher overcame this objection with a simplicity typical of mathematicians: Let X_i_ be the ith of n genotypes with frequency x_i_, then w_i_ is its fitness. On this basis, a discrete selection equation (the continuous one is not used here) can be formulated, which in the simplest case takes the form:

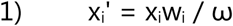

with

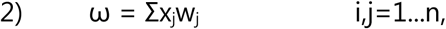

and Σx_i_=1. x_i_’ is the frequency of genotype X_i_ in the next generation and ω is the mean fitness. As Tiefenbrunner 1995^15^ has shown, if one considers the selection process to be fuzzy and the fitness values to be survival probabilities (rather than reproductive success), it is relatively easy to calculate the information change ΔH due to selection:

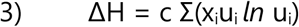

with

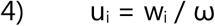

c is a constant. Since the problem remained that fitness is measurable only in exceptional cases, I published in 2020 another formulation for the change of the information by selection^16^:

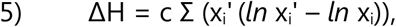

which has the advantage that fitness no longer occurs but only directly observable quantities (whose observation has recently become actually possible) are used. If recombination takes place, it can also be shown that the contributions to the information change Δh_j_ of the individual gene loci j can simply be added to the total change (g is the allele frequency and the index k runs over all alleles of the jth gene locus):

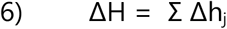

with

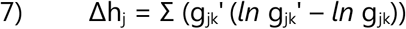

This finding is significant because, in general, the fitness of all gene loci does not simply sum to the total fitness of the individual.

Although it is possible to eliminate fitness, it would still be desirable if model examples were available in which all fitness values are always known during the entire selection process. Such examples can indeed be found, e. g. in the context of frequency-dependent selection, where the fitness of the phenotype depends on its frequency. In the past century, the chirality in snail shell coiling, has been well studied^11^. Snails can have a left- or a right-coiled shell and in an endemic species of a Pacific island, individuals with both shell types were observed in different valleys, but in the population of a valley dominated always either one or the other (the spezies is extinct in the wild since 1987). Since both shell types are presumably the same apart from chirality, one could assume that both were equally well adapted to the environment. Did selection take place here at all or was this a result of ‘random walk’? And in this rather special case, did selection also result in a quantitative change of information? These are the questions that will be dealt with in the following.

To analyze the claim that natural selection can also be of importance outside the animate nature, and also whether it could be a fundamental principle, I choose yet another example of frequency-dependent selection. H. Haken^9,10^ has described the emergence of laser light as a selection-analogous process of self-organization, but has not used the usual concepts of evolutionary biology. I want to make up for this here. Thus, the goal is to investigate the extent to which these concepts can be applied to a very simplified, laser-like, model for the generation of coherent light and to determine whether frequency-dependent selection in this framework causes an increase in information.

### Positive frequency-dependent selection in the snail *Partula suturalis*

The gastropod genus *Partula* includes species with different chirality, but most species have either only specimens with a left- or with a right-coiled shell. *P. suturalis*, now extinct in the wild, on the other hand, is polymorphic. Usually more or less monomorphic subpopulations occurred spatially separated on the island of Moorea. The local monomorphy was presumably the result of frequency-dependent selection and thus serves as an interesting example for the questions addressed here. It is unusually complex in that phenotype and genotype do not always coincide.

#### Inheritance

The mode of inheritance is relatively simple. A single gene locus determines the chirality, with the sinistral morph being inherited dominantly. However, the expression occurs with a delay of one generation, i. e. the maternal genotype and not the own defines the phenotype (*P. suturalis* is hermaphrodite, nevertheless, of course, the gametes are bimorphic). This circumstance imposes a significant complication, because it causes that almost any phenotype-genotype combination can be observed. If we denote the phenotypes with L and R, the diploid genotypes with ll, lr (=rl) and rr, then the following combinations are possible: L(ll), L(lr), L(rr), R(ll), R(lr) and R(rr). R(ll) does not occur de facto, but all the others do. If we denote by x the relative frequency, then obviously:

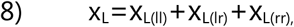

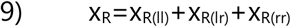

and x_R_+x_L_=1. Furthermore:

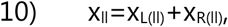

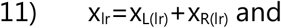

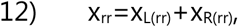

with x_ll_+x_lr_+x_rr_=1.

The haploid genotypes or allele frequencies, respectively, are calculated according to:

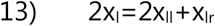

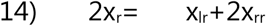

We are now interested in the frequencies of diploid genotypes and also phenotypes in the next generation, assuming, of course, that these are known in the present generation. If there were no selection, the population would remain in Hardy-Weinberg equilibrium. The allele frequencies could be calculated from Eq. 13 and Eq. 14, and the frequencies of the diploid genotypes in the next generation would be:

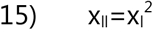

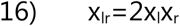

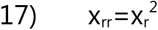

However, we assume that selection takes place and therefore the situation is far from simple. We have to look very closely at the mating event in particular. There are six different possibilities for recombination of the three diploid genotypes. Fig. 1 shows for all possible mating combinations the resulting output, i. e. the next generation.

**Fig. 1:**
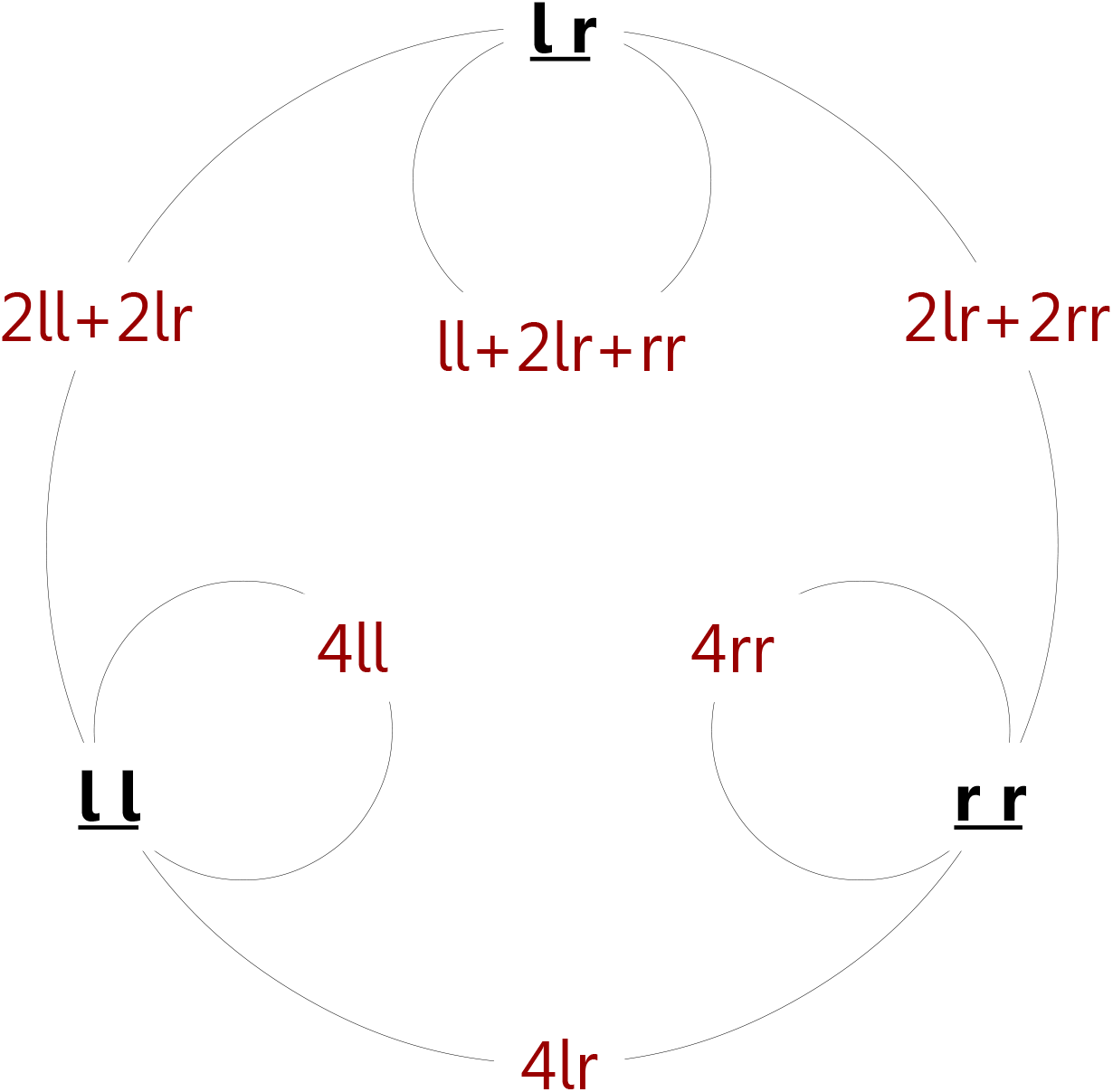
The figure shows the possible mating combinations and the genotypes of the recombinants (red) and their frequencies (the sum is four in each case). The graph represents the transition from one diploid generation to the next.

With the knowledge we now have, mating tables can be created, which we need to determine the frequencies of the diploid genotypes and also the phenotypes in the next generation. In addition to the information given in Fig. 1, these also include statements about the phenotype of the offspring. Crucially, therefore, we also need to consider which partner has the maternal (♀) role and which has the paternal (♂) role. Since each individual can be both, mother and father, the frequency is the same for both.

Accordingly, the frequencies of the phenotype-genotype combinations x_L(ll)_’ or x_R(ll)_’ (Table 1, left matrix) in the next generation, for example, would have to be determined by a two-step calculation:

**Table 1:**
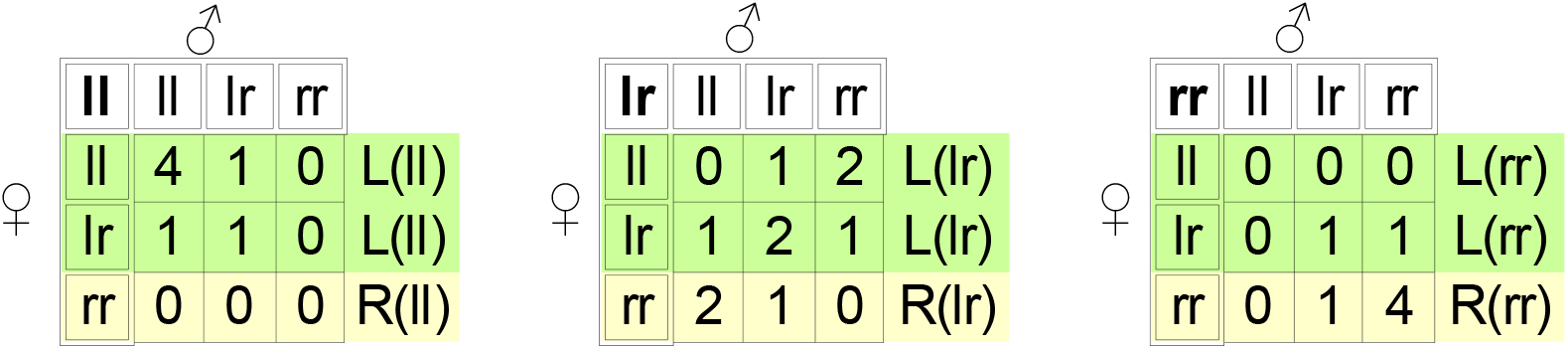
Generation transition matrices. Number of recombinants as a result of the combination (pairing) of two genotypes. Which one is the maternal one is crucial, as it alone determines the chirality of the offspring. Bold: recombinant genotype (next generation); leftmost column: maternal genotype; top row: paternal genotype. Maternal ll and lr determine a left-coiled shell in the offspring, rr a right-coiled one. Rightmost column: phenotype-genotype combination of the offspring.

**Table 2:** Fitness matrix for pairing between partners of the same and different chirality.

|  |  |  |  |
| --- | --- | --- | --- |
|  |  | ♂ |  |
|  |  | L | R |
| ♀ | L | 1 | 1-a |
|  | R | 1-a | 1 |

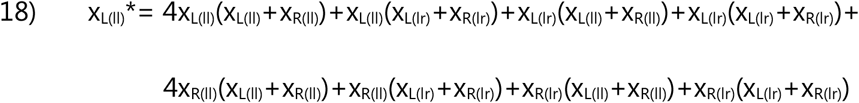

and

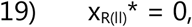

respectively. The remaining equations for the other phenotype-genotype combinations can be found in supplement 1. Eq. 18 could of course still be simplified because, e.g., x_L(ll)_+x_R(ll)_ = x_ll_ (Eq. 10), etc., but the form chosen for Eq. 18 will prove necessary in the next section. Finally, having calculated x* in this way for all phenotype-genotype combinations, we obtain:

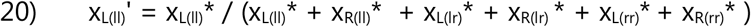

and

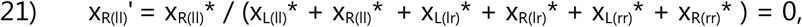

so that finally x_L(ll)_’+x_L(lr)_’+x_L(rr)_ ‘+x_R(ll)_’+x_R(lr)_’+x_R(rr)_’=1. However, things would only be that easy if there wouldn’t be selection.

#### Selection

We assume that chirality per se does not cause a fitness difference, i. e., a left-coiled shell should be equally good as a right-coiled one. In a series of laboratory experiments, Johnson found in 1982^11^ that the cause of frequency-dependent selection in *Partula suturalis* is lower mating efficiency as a consequence of the morphological differences when the bodies of the mates have different chirality. We therefore obtain the following symmetric matrix for mating (0≤**a**≤1):

If the mating partners have different chirality, the relative frequency of the offspring is reduced by **a** (tab. 2). **a** is 0.28 in the laboratory, but certainly reached higher values in the field. If selection is taken into account, e. g. Eq. 18 according to Tab. 2 takes a different form:

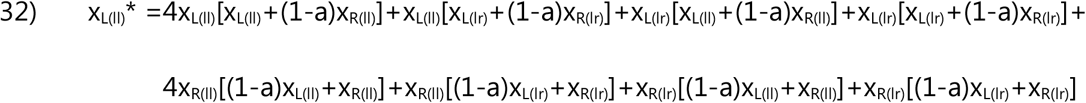

It cannot be seen immediately from the Eq. 32 and the other equations in supplement 1 that a frequency-dependent selection takes place and thus a simulation is necessary. This leads, because the two phenotypes can be combined with almost any genotype, only to a slow change of the geno- and phenotype frequencies. To speed it up somewhat, I chose **a**=0.8 in the simulation to be discussed below, which is possible without limiting the generality of the conclusions (Fig. 7). The initial conditions were set as follows: A value for x_l_ is determined (and thus also for x_r_=1-x_l_), the population is initially in Hardy-Weinberg equilibrium (Eqs. 25-27), and furthermore, in the first generation the phenotype corresponds to the genotype (thus, only three of the potential six combinations are present). We ask for the information change due to selection, which is calculated according to Eq. 5 for the frequencies of diploid genotypes, phenotypes, and phenotype-genotype combinations. The information change is shown only from the third generation onward because the chosen initial frequencies of the phenotype-genotype combinations would not normally occur if the phenotypes were inherited maternally.

The simulation was performed over 100 generations for two initial conditions (Fig. 2): if initially x_l_=0.5, allele variant l and phenotype L eventually prevail, but complete displacement is not observed in the selected time interval. On the other hand, if x_l_=0.3 applies initially, the frequency of allele r increases and also of phenotype R, but they do not fully prevail in the chosen time interval. The information content increases due to frequency-dependent selection in both cases studied. However, the calculation must be done using the frequency changes of the phenotype-genotype combinations, since considering only the genotype or the phenotype would result in too low values for the information change. This is a consequence of the fact that in *Partula suturalis-*individuals the genotype and phenotype do not correspond to each other.

**Fig. 2:**
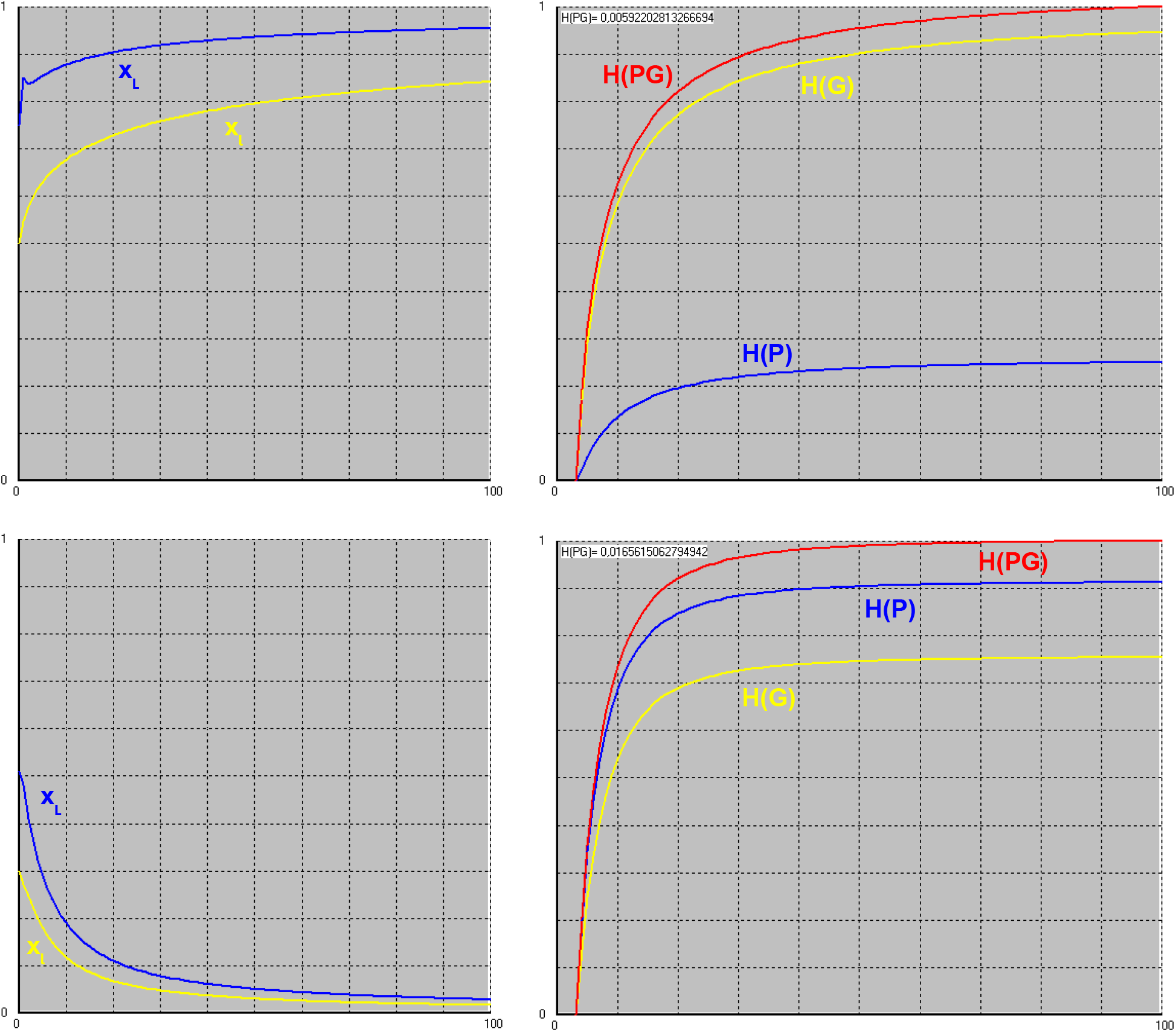
Density-dependent selection in the Partula suturalis-model with initial conditions x_l_=0.5 (top) and x_l_=0.3 (bottom). In each case, the left figure shows the relative frequency for allele l or phenotype L in the population, whereas the right figure shows the cumulative sum of the information change ΔH up to the current generation relative to the sum over all generations when the information change is calculated from the frequencies of diploid genotypes H(G), phenotypes H(P), and phenotype-genotype combinations H(PG) according to Eq. 5. For Partula suturalis, where genotypes and phenotypes do not correspond to each other, only the last approach is correct.

In the model, dextral and sinistral morphs were assumed to have equal fitness. For the sake of completeness, it should be mentioned that this was not always the case for *Partula suturalis* in the wild, which could be seen in the distribution of the dextral populations^11^. Namely, they were observed where other species of the genus *Partula* (now extinct or extinct in the wild, too) occurred that were sinistral. The reason is likely that mating happened between individuals of different species, but the offspring were then infertile. Inter species hybridization, however, occurred less frequently and was less successful when chirality differed, which was advantageous in this case. In the model, inter species interactions as they happened in reality were ignored, so in this framework the assumption of equal fitness for both morphs seems justified.

We note that frequency-dependent selection leads to information increase and now turn to the next, the non-biological (and more speculative) model.

### Positive frequency-dependent selection and information change in the laser-like model

Since “frequency” has a special meaning in physics, in the following chapter I will no longer speak of “frequency-dependent” but instead of “density-dependent” to avoid misunderstandings.

Laser light has extraordinary properties: it is monochromatic, monoaxial and monophasic. In its formation, the usual ingredients of evolution, namely exponential propagation, selection by resource limitation – and even inheritance and mutation – play a significant role. In the following, we want to identify them and design a laser-like model in which the information change during the selection process can also be described.

Laser light is generated far from thermodynamic equilibrium in an active medium where atoms or molecules are in an excited state (in which they must be artificially maintained) so that photons can easily be emitted. A. Einstein discovered in 1917^5^ that in addition to the spontaneous emission of photons, there also exists an induced one (he called it „Zustandsänderung durch Einstrahlung”), i. e., photons can excite the emission of others with exactly the same properties, resulting in a kind of chain reaction. The ‘explanation’ of this phenomenon was later given by quantum theory and has something to do with the property of all bosons (not only photons) to prefer to be in the same state^6^. For photons it follows: The probability that an atom emits a photon in a certain state increases by a factor of m+1 if there are already m photons in that state ^7^. Something very similar is observed, for example, in the population of a species of parthenogenetic animals: the probability of a birth (or a hatching) in a given period increases with the number of individuals m. The difference between the factors m and m+1 is particularly relevant when m=0. In the quantum world – contrary to the biological one – ‘primordial generation’ occurs but here is called ‘spontaneous emission’. The potential for exponential propagation therefore exists, also some kind of heredity, but how does selection occur?

The induced emission is the more successful the longer the light remains in the active medium which is bordered by two parallel mirrors, and the light with the wavelength that fits in exactly and also has the right direction of propagation is preferred in this respect and thus is fitter. But what about the phases? Aren’t all of them equally fit? Initially, this is indeed the case.

In the laser-like model in the context of density-dependent selection, we are only interested in phase selection. So we assume that the selection on wavelength and propagation direction has already been done, but that the phase is initially equally distributed, which results in an amplitude of zero (the amplitude square corresponds to the light intensity, i.e. no photon can be detected). In contrast to reality, where the phase difference between two photons can be arbitrarily small, we allow only discrete differences. There are n ‘phase classes’ (thus we have our genotypes) which differ from each other by an integer multiple of a phase angle α=2π/n (fig. 3).

**Fig. 3:**
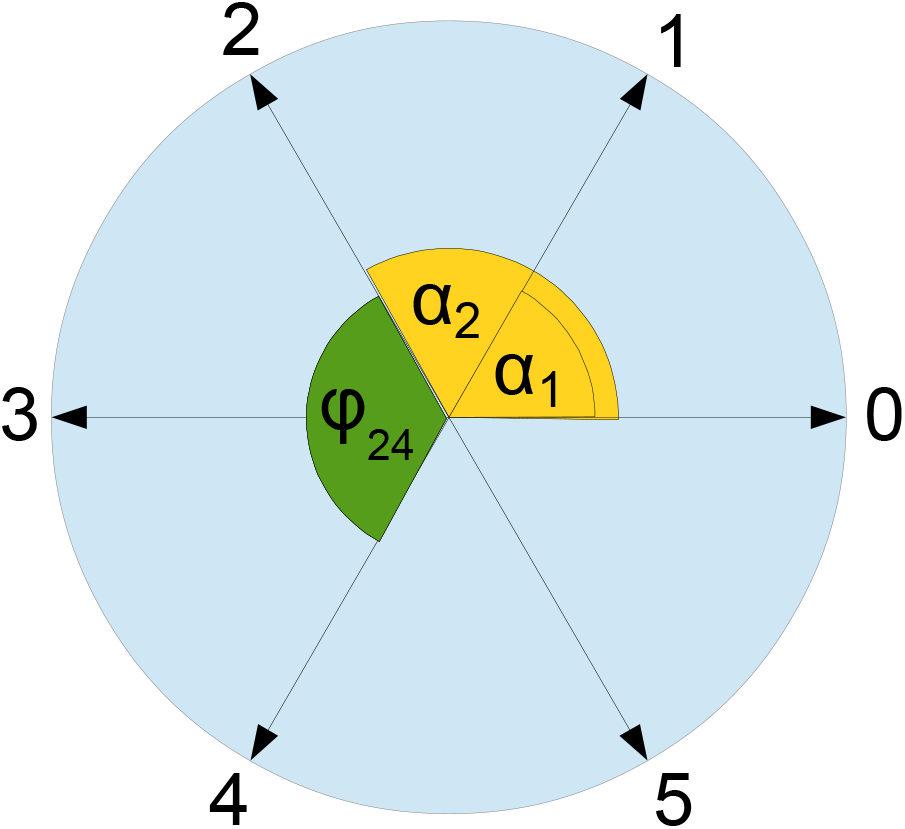
The photons of the model oscillate in different phases, where the phase angle is an integer multiple of α=2π/n. Here n=6.

Let n be an even integer. Let the ith phase class be X_i_ with 0 ≤ i ≤ n-1 and let the relative frequency of photons of X_i_ be x_i_. Then

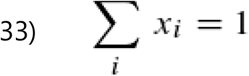

Furthermore, according to Fig. 3:

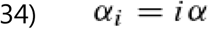

and

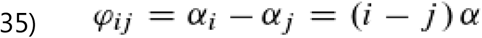

where 0 ≤ i,j ≤ n-1.

As in the previous model, we now need a hypothesis about the transition from one photon-’generation’ to the next. We have two sources of information for this: on the one hand, we know that photons have a tendency to want to be indistinguishable. On the other hand, we know how interference takes place. Using this, we postulate a kind of recombination. The naive assumption is that whenever a X_i_ - photon “meets” (whatever this may mean) a X_j_ - photon, an event which occurs with likelihood x_i_x_j_, there is a transition X_i_↔X_j_ with probability P(X_i_↔X_j_) that leaves both in the same state (Fig. 4). The process is supposed to be symmetric: P(X_i_→X_j_)=P(X_i_←X_j_) so that we use P(X_i_↔X_j_) for both; the other reason to do so is that the recombinant photons are not either or but they are both, a phenomenon called ‘superposition’. This kind of intermediate inheritance is of course unknown in animate nature.

**Fig 4:**
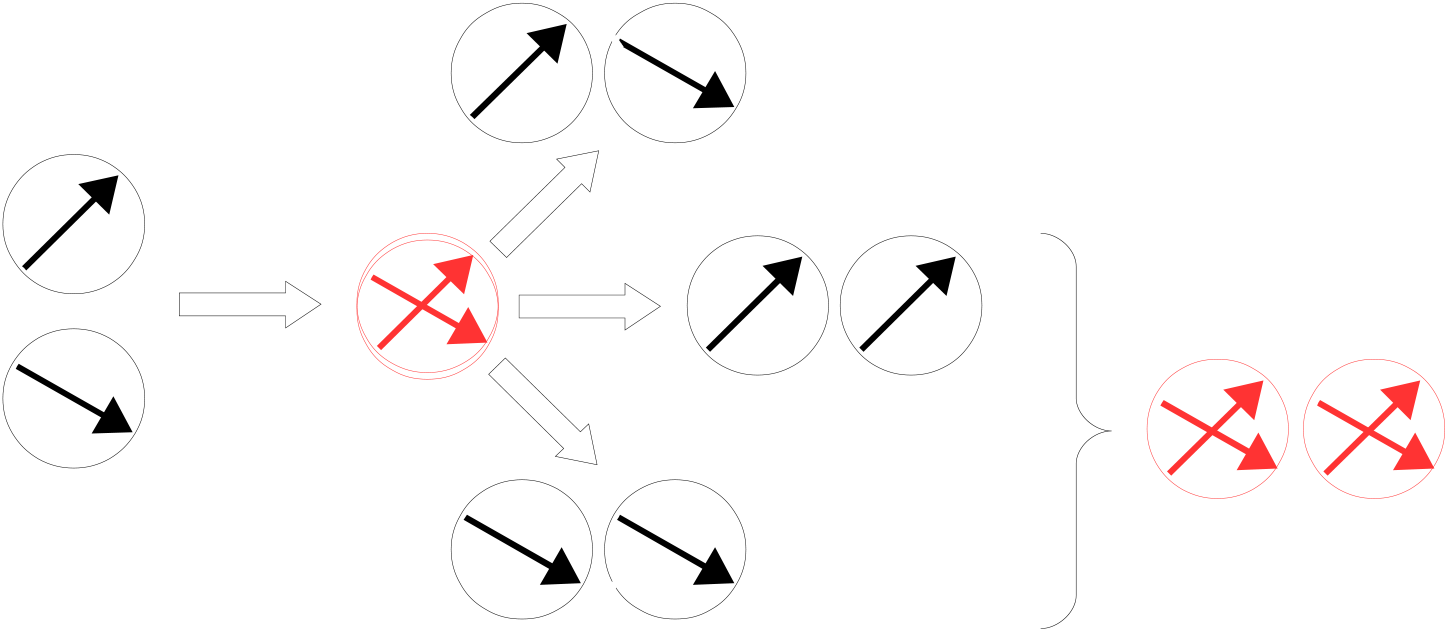
If two photons (circles with arrow) “meet”, a kind of recombination may occur with transition probability P(X_i_↔X_j_) that leaves both in the same state, either in the one or the other (alternatively we may say they are in superposition). This is a simple and naive assumption that must be justified by the results.

Why can’t the result be photons with a phase that lies between those of the initial ones? Simply because this phase could then lie between the phase classes allowed in the model and because it does not fit to the phenomenon of interference. In order to determine the fitness of the X_i_-photons, we need to know the transition probability P(X_i_↔X_j_), which we try to guess from the theory of wave interference: The resulting length of the amplitude a_ij_ is obtained from the both individual amplitudes a_i_ and a_j_ according to Fig. 5, where φ_ij_ is the phase angle between the phase classes X_i_ and X_j_.

**Fig. 5:**
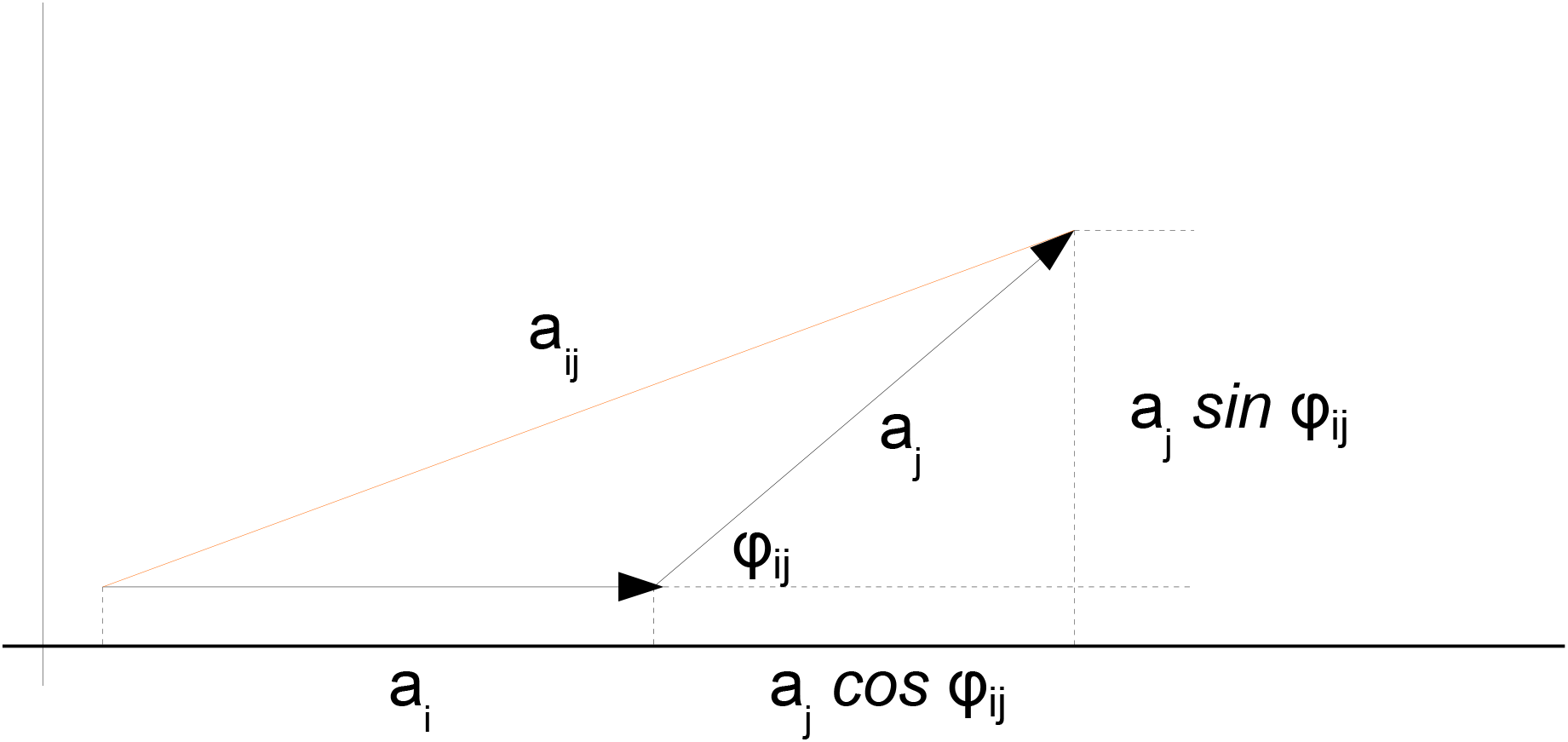
Interference of two waves of coherent light of the same frequency but different phase; amplitude addition.

So it follows from the Pythagorean theorem:

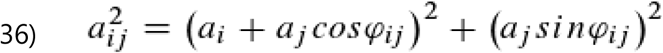

after some calculations using sin^2^φ + cos^2^φ = 1 we get:

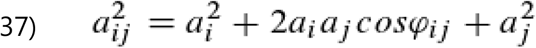

Since P(X_i_↔X_j_) is a probability it can have only values between zero and one, we choose

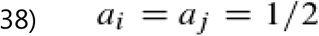

so that

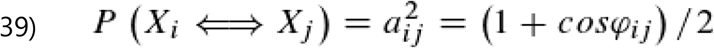

Later, however, it will turn out that the factor ½ gets canceled out, so that finally we use

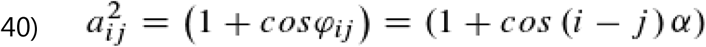

instead. From this, the change of the frequencies x_i_ within ‘one generation’, as well as the fitness values w_i_ and the mean fitness ω of the ‘photon population’ can now be calculated. According to R. A. Fisher^8^, the following applies in the discrete selection model (see also equations 1 and 2):

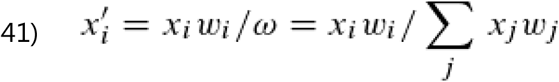

where x_i_’ is the relative frequency in the next generation and based on our considerations it follows that

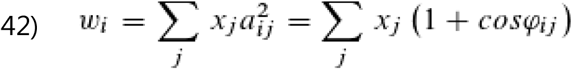

For mean fitness ω, we find of course:

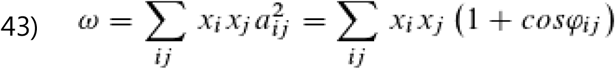

This gives us everything we need to investigate with an example what happens if we assume that initially x_i_ = 1/n for all x_i_. Nothing happens at all. But if we increase, say, x_0_ by a tiny amount ε and decrease all the others so that they are equal to each other, exactly so that Σ x_i_ = 1, then something happens (Fig. 6).

**Fig. 6:**
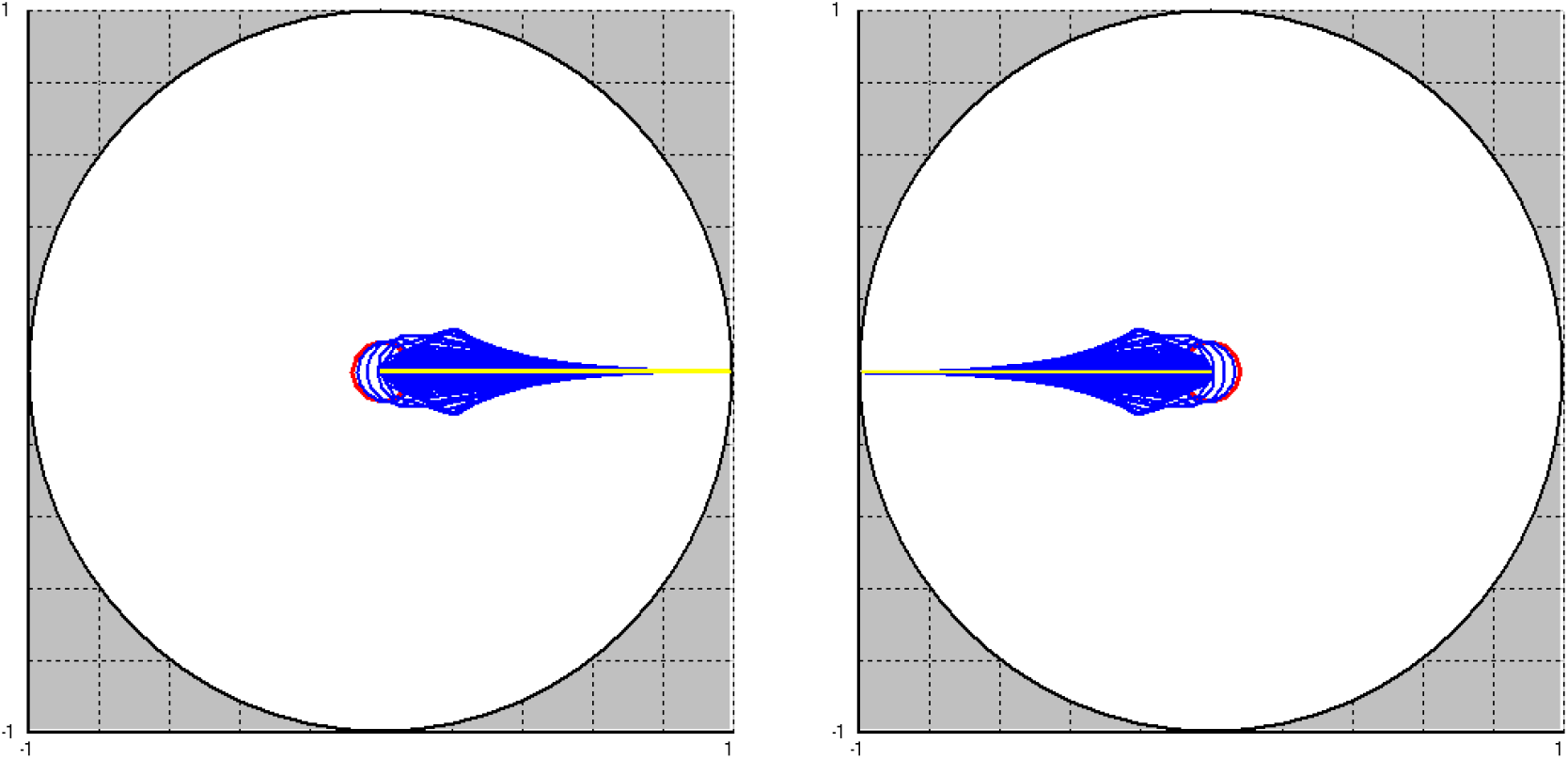
Change of the ‘photon population’ in the laser-like system by density-dependent selection of the phase. In the left image, x_0_ was assumed to be more abundant than the other phase classes by ε=0.0001 (n=12). On the right, ε=-0.0001. Red: initial state, yellow: final state. All other generations were drawn in blue. Every third generation was plotted. The simulation ran for 102 generations.

Density-dependent selection causes the smallest initial advantage to be sufficient for a particular phase to fully prevail (Fig. 6, left). If only one phase has an initial disadvantage, the phase shifted by π prevails (Fig. 6, right). For the latter, of course, there is no pedant in the realm of biology.

For the examples, we can also calculate some important parameters during the simulation run, namely the mean fitness ω, the square of the total amplitude A^2^, and finally also the information change ΔH per ‘generation’, which is what we are actually concerned with. For the latter we can use the equation (see also equation 5)

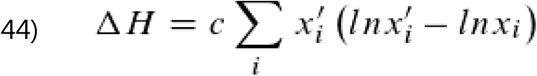

from ref. 16 to calculate the change of H. For the constant c we choose the value one. Fig. 7 shows how the total amplitude can be determined.

**Fig. 7:**
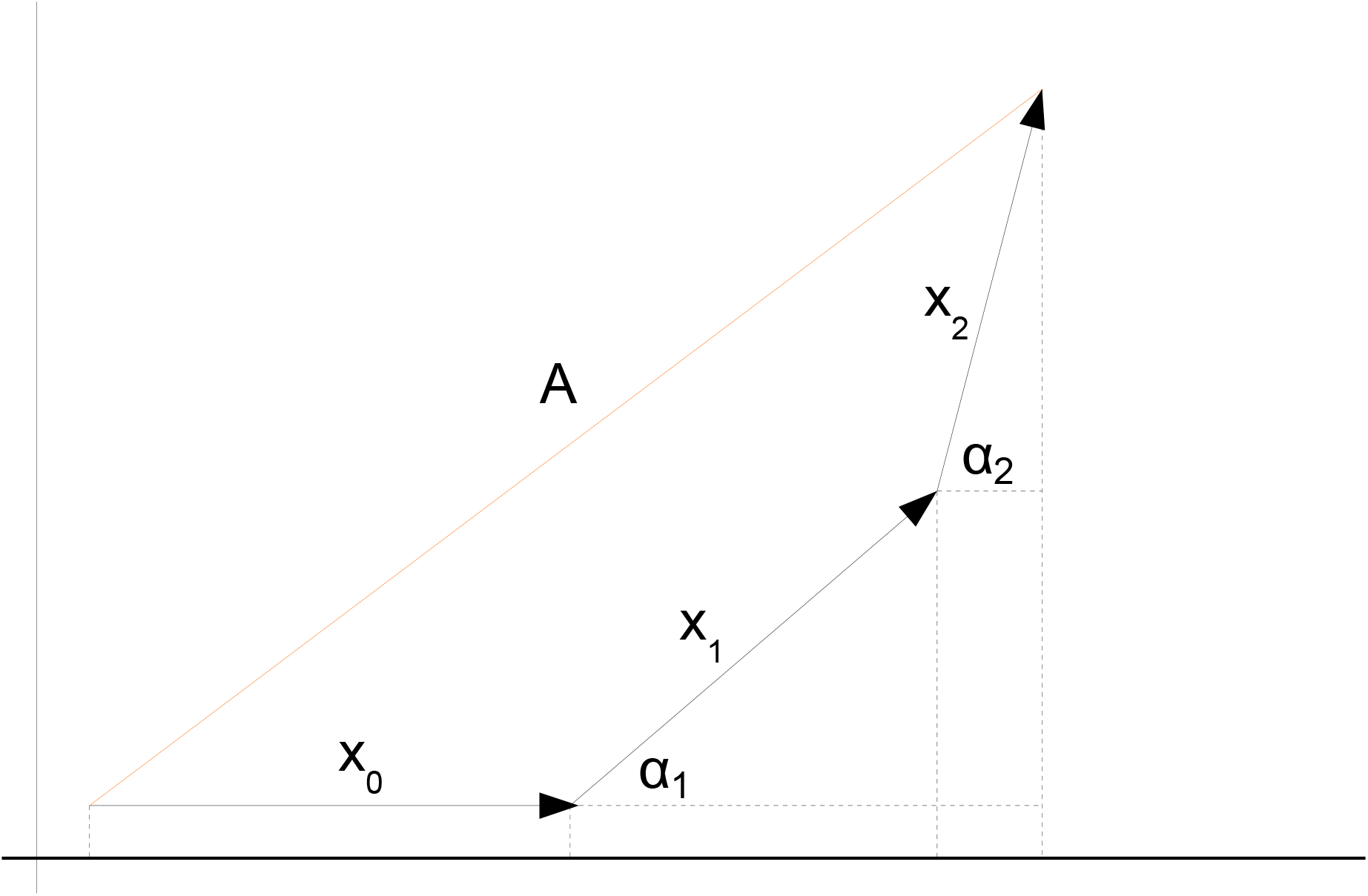
Calculation of the length of the total amplitude (resultant) by lining up arrows (adding amplitudes).

As one can see from the figure, using the Pythagorean theorem gives:

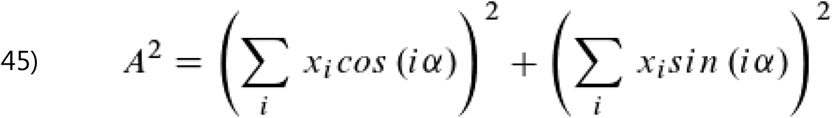

Alternatively we may write:

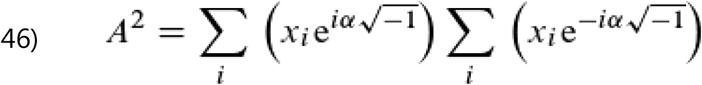

Thus, all components for a comparison are available (Fig. 8).

**Fig. 8:**
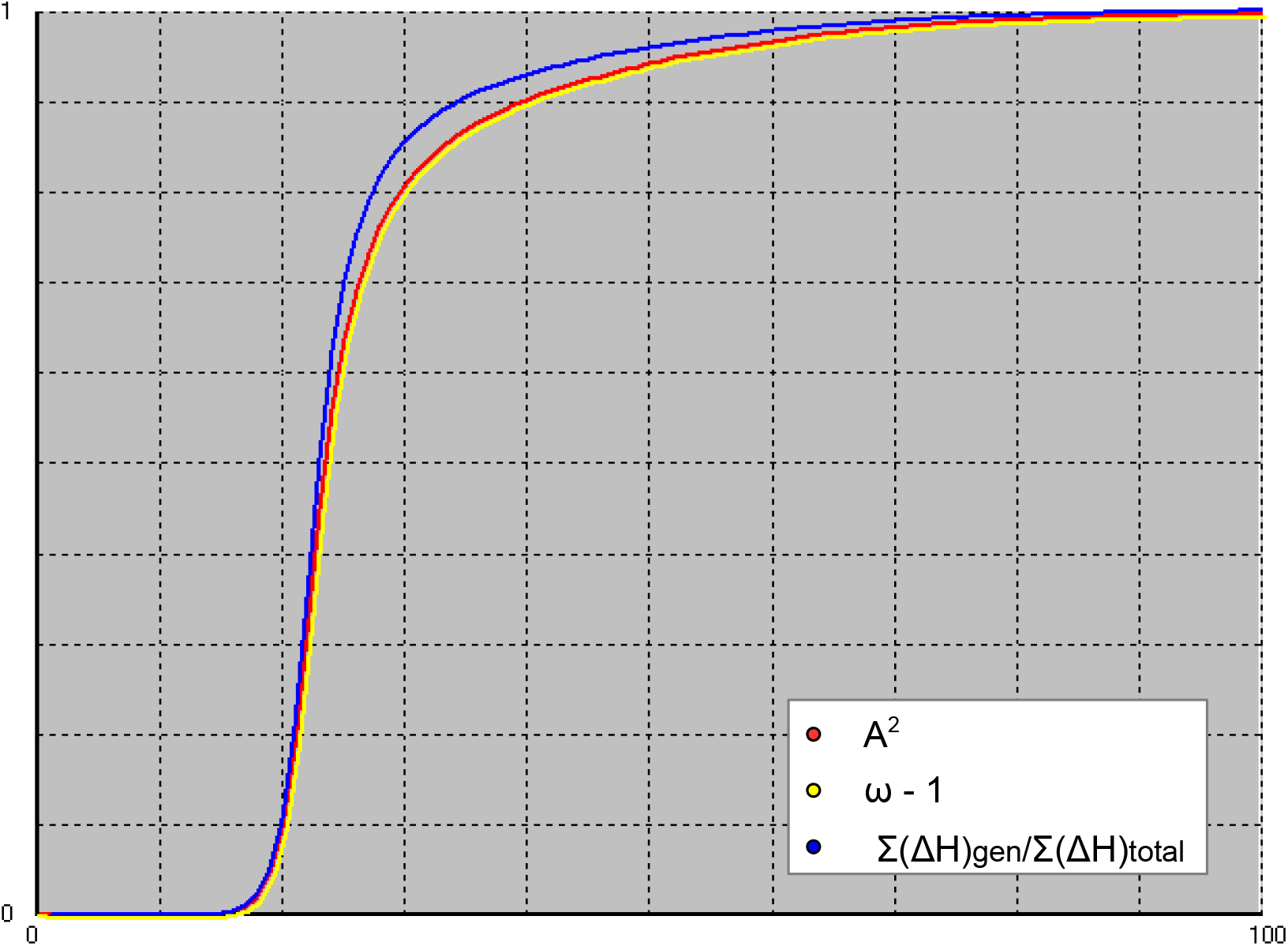
Change of the ‘photon population’ in the laser-like system by density-dependent selection of the phase. For n=12 and 100 generations the square of the total amplitude A^2^, the mean fitness decremented by one (ω-1) and the sum of the information change ΔH up to the current generation relative to the sum over all generations (Σ(ΔH)_total_=0.1788 ‘selbit’ per photon) are shown. The yellow and red curves are drawn slightly offset from each other, since one would otherwise completely overlap the other.

In Fig. 8, it is immediately noticeable that the yellow (ω-1) and red (A^2^) curves overlap completely and are therefore identical. This does not change if the number of phase classes n or the number of generations is changed. It is therefore obviously valid:

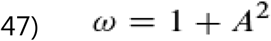

as long as Σ x_i_ = 1. Eq. 47 is very interesting and hence I will give a general proof in supplement 2. So there are different mathematical ways to calculate the total amplitude; besides Eq. 45, we could also use complex numbers (Eq. 46), or just Eq. 43 in connection with Eq. 47. At first, there is nothing more behind it. But in the model of phase selection it becomes something new, namely that behind the total amplitude there is a transition matrix with transition probabilities corresponding to the mean fitness – an unexpected meeting point of two seemingly completely different theories. We can now use the relationship described by Eq. 47 to calculate the change in information. The equation for information change by selection can also be represented this way (see also equation 3):

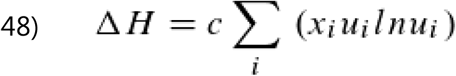

(ref. 15) with

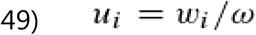

So by substituting we finally get:

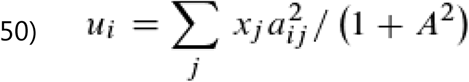

Because 0 ≤ A^2^ ≤ 1, it follows that 0 ≤ ln(1+A^2^) ≤ ln2.

In the context of selection, an increase of the mean fitness always corresponds to an increase of the information content and since within this model the amplitude square practically takes over the role of the mean fitness, the question arises whether – if we assume a constant number of photons – a larger amplitude generally can also be attributed to a higher information content (lower entropy). This assumption may have its justification. For example, in the focal point of a concave mirror the phases of the photons – imagined as arrows – all point in the same direction and the intensity of light is high, at other places, however, they are more or less confused. The situation is somewhat similar to another one we know: If the ‘directional arrows’ of the movement of air molecules are randomly distributed and thus the entropy is high, we do not perceive any air movement, but we do if most of them point in one direction (low entropy). Then we feel a wind or even a storm.

However, since our model is very simplistic (in reality there are no discrete time steps or phase classes and the equation for the information change has been developed from the classical information theory) it would certainly not be justified to claim physical (beyond model) significance for it. Further more or less obvious physical consequences from the model will therefore not be discussed here. However, we conclude that density-dependent selection of the phase leads to an increase in information.

For both models (laser-like and *Partula suturalis*), there are quantitative statements on the change in information content per individual. Is it therefore possible to compare the models in this respect? One must not forget that Eq. 12 has a constant c for which the value one was arbitrarily chosen in each case. In reality, however, the value for c in both models is certainly very different (and currently unknown). A statement such as: “The information change per individual or photon is larger in the laser-like model, because H is also larger”, would therefore be wrong.

If one wants to make such models comparable and also make physically relevant statements, theoretical investigations will not be sufficient. Experiments will also have to be carried out. To start with, model organisms with a very short generation sequence are needed, but which reproduce sexually so that recombination takes place in each generation. Then it will be possible to study the influence of physical conditions, such as temperature on selection, and also that of other environmental impacts, but also how the number of genes affects or whether there is an upper limit for information acquisition per unit time, what effect selfish genes or cooperation have. The theory of information acquisition by selection could therefore open up new research opportunities.

## Supplement 1

In this supplement all transition equations for the *Partula suturalis*-model are presented:

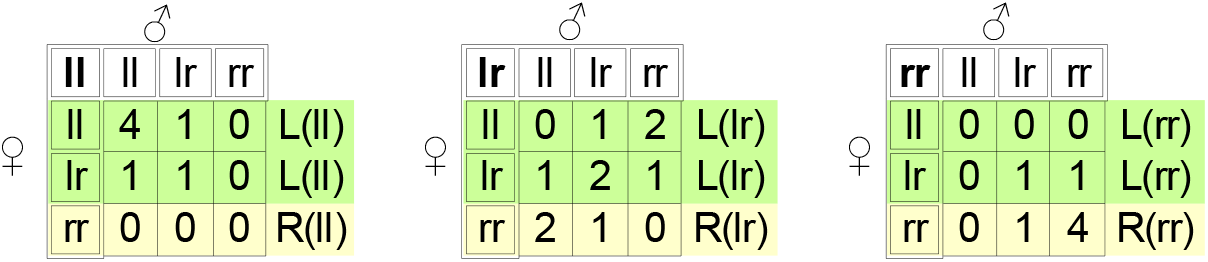

From the left matrix:

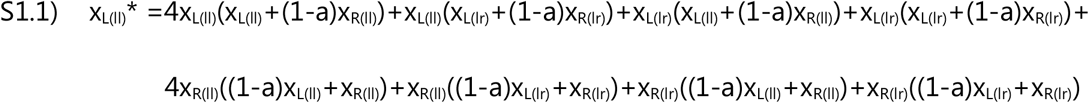

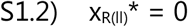

From the matrix in the middle

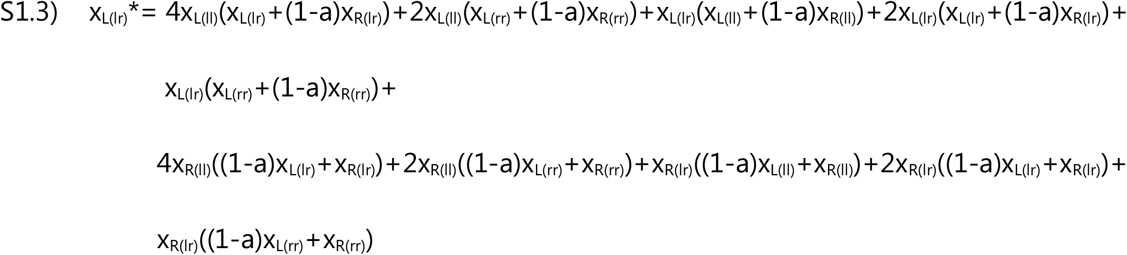

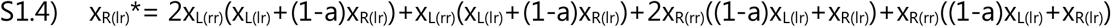

From the right matrix

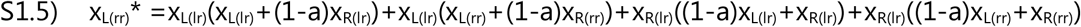

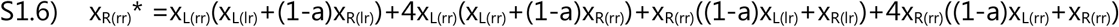

### Supplement 2

We want to show that Eq. S2.1 (Eq. 47) is valid

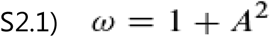

if (from Eq. 43)

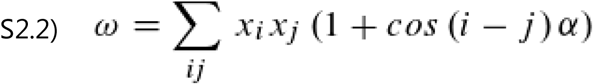

and (see also Eq. 46)

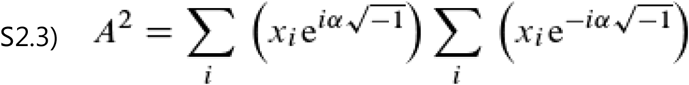

From Eq. S.2.2 it follows:

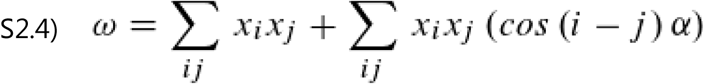

and thus

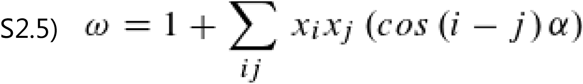

because Σx_i_x_j_=1 if Σx_i_=1. From S2.1 and S2.5 it follows:

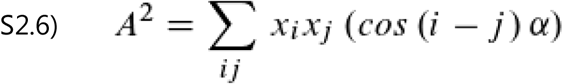

Let us define b_ij_^2^:

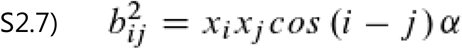

so that A^2^=Σb_ij_^2^. Now we calculate the sum of b_ij_^2^ and b_ji_^2^:

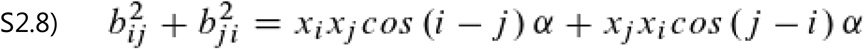

Since cos (-γ) = cos γ it follows:

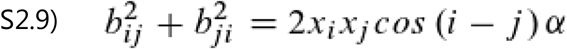

And because cos 0 = 1, from S2.7 follows: b_ii_^2^=x_i_^2^.

Now let’s start with Eq. S2.3, which can also be written like this:

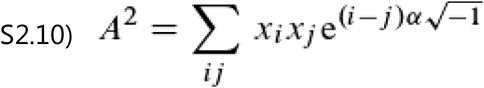

and

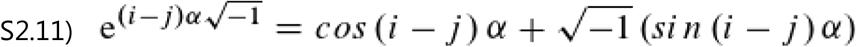

We may define a_ij_^2^:

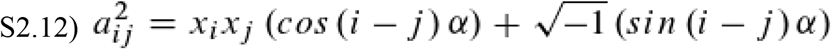

so that A^2^=Σa_ij_^2^. Now let us calculate the sum of the pair a_ij_^2^ and a_ji_^2^:

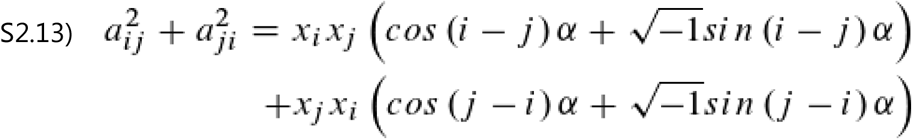

Because of sin (-γ) = - sin γ

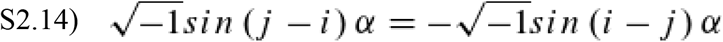

we finally get:

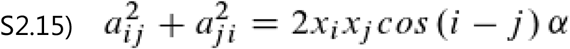

Thus we see that

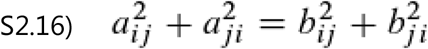

and furthermore that

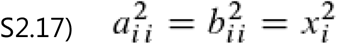

and hence that Eq. S2.1 is valid.

